# *In situ* miRNA delivery from a hydrogel promotes osteogenesis of encapsulated mesenchymal stromal cells

**DOI:** 10.1101/712042

**Authors:** James. Carthew, Surakshya. Shrestha, John. S. Forsythe, Ilze. Donderwinkel, Vinh. X. Truong, Jessica. E. Frith

## Abstract

Hydrogels have many properties that emulate biological tissues and are therefore attractive candidates for use in tissue engineering. In particular the encapsulation and subsequent differentiation of mesenchymal stem/stromal cells (MSCs) is a strategy that holds great promise for the repair and regeneration of bone and cartilage. However, MSCs are well-known for their sensitivity to mechanical cues, particularly substrate stiffness, and so the inherent softness of hydrogels is poorly matched to the mechanical cues that drive efficient osteogenesis. This limits the success of bone tissue engineering using MSCs encapsulated in a hydrogel. One approach to overcome this limitation is to harness mechanotransductive signalling pathways and override the signals cells receive from their environment. Previous reports have shown that the mechanosensitive miRNAs, miR-100-5p and miR-143-3p can enhance MSC osteogenesis, but this required a complex multi-step procedure to transfect, encapsulate and differentiate the cells. In this study, we develop and characterise a facile system for *in situ* transfection of MSCs encapsulated within a light-crosslinkable gelatin-PEG hydrogel. Comparing the influence of different transfection agents and hydrogel compositions, we determine the factors affecting transfection agent release and MSC transfection, showing that it is possible to transfect MSCs with miRNAs *in situ*. We then compare the efficacy of both pre-transfection and *in situ* transfection on the osteogenic capacity of hydrogel-encapsulated MSCs, demonstrating superior mineralisation and osteogenic gene expression for *in situ* transfected samples. Our platform therefore demonstrates a simple, one-pot system for delivery of pro-osteogenic miRNAs and *in situ* transfection that is able to enhance MSC osteogenic potential without the need of multi-step transfection procedures, thus demonstrating significant promise for bone tissue engineering.

## 1 Introduction

For many years, the application of mesenchymal stem/stromal cells (MSCs) has been studied as a means to enhance regeneration of bone *in vivo*, through either direct injection to the site of injury, or when seeded onto ceramic implants for larger fracture areas [1, 2, 3, 4]. One drawback of these theraputic approaches is the inherant low cell viability following stem cell transplantation, attributed to the host immune response, high shear forces associated with the injection process and mechanical shear from the surrounding microenvironment. As a result, hydrogels have attracted great interest as a way to protect cells during delivery and provide a favourable microenvironment to support osteogenesis thereafter. Through a combination of their compositional versatility and high degree of biocompatibility, many hydrogel formulations demonstrate minimal cell mortality and functional bone regeneration both *in vitro* and *in vivo* following cell encapsulation [5, 6, 7, 8, 9]. Their innate ability to take on the shape of complex defects makes them uniquely suited for bone tissue engineering, however one critical drawback, particularly for MSC encapsulation is their inherently soft nature. Due to their high water content, injectable hydrogels have a significantly lower modulus than that of surrounding bone tissue. Taking into account the high degree of mechanosensitivity shown by MSCs, coupled with extensive literature demonstrating the impact of substrate stiffness in directing stem cell lineage (with stiff substrates (40 – 70 kPa) favouring osteogenic lineage and soft substrates (0.5 – 5 kPa) favouring adipogenic lineage) [10], the use of these systems for bone regeneration is likely sub-optimal.

One strategy to improve the osteogenesis of encapsulated MSCs is to deliver drugs or biomolecules that act upon mechanotransductive signalling mechanisms and over-ride the physical cues that the cells derive from the soft hydrogels. In this manner it may be possible to artificially induce the MSCs to respond as if they were on a high-modulus, pro-osteogenic substrate. Our previous work identified a panel of mechanosensitive microRNAs (miRNAs) and showed that they could be used to modulate MSC osteogenesis and improve bone formation in MSCs exposed to a soft substrate [11]. Other studies have since further demonstrated the osteoinductive nature of miRNAs [12, 13]. miRNAs are a class of endogenous small, non-coding RNAs that positively or negatively regulate gene expression and cellular processes via RNA interference [14, 15, 16]. By targeting mRNA transcripts post-translationally, they provoke either translational repression or degradation, depending on the degree of sequence homology, and thus play critical roles in the regulation of cell signalling pathways. The use of miRNAs to drive cell fate and enhance tissue regeneration is of great interest due to a combination of their powerful effects on biological process, relative ease of delivery and inexpensive nature [17]. miRNAs are therefore of significant interest in tissue-engineering and hold significant promise to optimise regenerative outcomes.

Our previous study showed that MSCs transfected with the pro-osteogenic miRNAs, miR-100-5p and miR-143-3p had increased osteogenic differentiation potential and that this was mediated by modulation of mTOR signalling [11]. One drawback was that this study relied upon the pre-transfection of MSCs with osteogenic miRNAs prior to encapsulation. Although cells were successfully transfected, this multi-step procedure was a cumbersome process requiring repeated cycles of media changes and manipulation during the transfection and subsequent encapsulation. To overcome this, here we develop a system to facilitate *in situ* transfection of MSCs encapsulated in a hydrogel, significantly simplifying the process to a one-step procedure that encapsulates both MSCs and miRNAs complexed with transfection agents, at the moment of hydrogel crosslinking.

We have developed and characterised a facile platform in which a polyethylene glycol (PEG) / gelatin-norbornene (GelNB-PEG) hydrogel, pro-osteogenic miRNAs and transfection agents are combined to enhance osteogenic differentiation of encapsulated MSCs *in situ*. Successful development of this system not only demonstrates the ability of miRNAs to direct MSC fate irrespective of microenvironment, but also takes steps towards the future translation of such systems for bone tissue engineering.

## 2 Materials and Methods

### 2.1 Synthesis of Poly(ethylene glycol) dithiol (PEGdiSH)

PEGdiSH synthesis was conducted in a one-step process. Briefly, 10 g PEG-2armOH (Mn = 2000 Da, 0.1 M, UK) and 9.5526 g of mercaptopropioinic acid (0.09 M, USA) were dissolved in 100 mL toluene, and two drops of H_2_SO_4_ (98 %) were added. The resulting solution was then heated to reflux at 120 °C under Dean-Stark conditions for 20 h. Toluene was subsequently evaporated off *in vacuo*, with the resulting oil dissolved in dichloromethane (100 mL) and then washed in saturated NaHCO_3_ solution (50 mL). The organic phase was dried over anhydrous magnesium sulfate (MgSO_4_), filtered, and concentrated to ca. 5 mL. The polymer was then purified by precipitation into diethyl ether.

### 2.2 Synthesis of Gelatin Norbornene (GelNB)

GelNB was synthesised based on previously reported methods [18]. Briefly, 1.00 g of 5-norbornene-2-carboxylic acid [7.20 mM] (Sigma, mixture of endo/exo) was dissolved in 20 mL dichloromethane, followed by addition of 1.12 g N-hydroxysuccinimide [9.72 mM] and 1.79 g of N-(3-(dimethylamino)propyl)-N’-ethylcarbodiimide hydrochloride [9.36 mM] (Sigma). The resulting solution was mixed at room temperature (RT °C) for 20 h.

The reaction mixture was then washed with 40 mL saturated NaHCO_3_ solution, twice with H_2_O, and dried over anhydrous magnesium sulfate (MgSO_4_). The dried solution was then evaporated *in vacuo*, yielding a white solid product (1.45 g, ca. 86 %). Next, 1.00 g gelatin powder (Sigma, porcine skin Type A) was dissolved in 20 mL N,N-dimethylformamide (DMF)/H_2_O (1:1 v/v) solution and stirred until completely dissolved. To this solution, the above resultant (5-norbornene-2-carboxylic acid NHS, 59 mg, 0.25 mM) dissolved in DMF (5 mL) was added with an additional 50 μL N,N-diisopropylethylamine (DIPEA) (Auspep, Australia). The resulting solution was left to stir at ambient temperature for 8 h, followed by dialysis in excessive deionised H_2_SO_4_ for 3 days in 3.5 kDa cut-off dialysis tubing. The purified solution was filtered using a 0.2 μm syringe filter, and subsequently lyophilized.

### 2.3 Hydrogel Polymerisation

Polymers (GelNB and PEGdiSH) and lithium phenyl-2,4,6-trimethylbenzoylphosphinate (LAP) photo-initiator were dissolved in Dulbecco’s phosphate-buffered saline (DPBS) (Life Technologies). A 200 μL mixture was prepared by mixing the corresponding amounts of stock solutions and DPBS to deliver two final solutions of 1 % (w/v) PEGdiSH, and 0.03 % (w/v) LAP with either 10 % (w/v) or 4 % (w/v) GelNB. Following stabilisation for 2 min, photo-polymerisation using an OmniCure S2000 (Lumen Dynamics) was performed for 10 min using a 400-500 nm filter at a calibrated light intensity of 10 mW cm^-2^. For MSC encapsulation studies, cell suspensions [2.5 million cells/mL] were created using the GelNB solution prior to addition of PEGdiSH and LAP components. Gels were then polymerised as described, and cultured at 37 °C for experimental duration.

### 2.4 Rheological Analysis

Rheological analysis was performed using a parallel plate strain-controlled rheometer (Anton Paar Physica). A top plate diameter of 25 mm was selected with a gap size of 0.5 mm, constant strain of 0.2 % and frequency of 1 Hz. Hydrogels were produced as stated above, with the addition of a 24 h swelling step conducted in DMEM supplemented with 10 % (v/v) fetal bovine serum (FBS) (Scientifix) at 37 °C.

### 2.5 Swelling and Degradation Measurements

Hydrogel precursors and component parts were prepared as described previously. The resulting mixture of gel stock solutions were pipetted into a 10 mm micro-well culture dish (ThermoFisher). For each sample, 200 μL droplets were formed, with the macromer mass used in the precursor recorded as *mp*. Gels were swollen in excessive DPBS for 24 h to reach equilibrium swelling, following which, gel mass was recorded as *ms*. Gels were subsequently vacuum-dried with the dry mass being recorded as *md*. The swelling ratio was calculated as *ms / md*, the equilibrium water content (%) was calculated as (*ms* - *md*) / *ms* x 100 %, and the gel fraction was determined as *md* / *mp* x 100 %. For the degradation measurements, hydrogels were incubated in DPBS at 37 °C, removed from solution and their mass recorded as *mt* every day. The degradation was measured as the percentage of *mt* to the mass of as gelled hydrogel at day 0 (*mt0*) as (*mt / mt0*) x 100 in triplicate.

### 2.6 Release Kinetics Analysis

To assess release kinetics of miRNAs from both 10 % and 4 % (w/v) GelNB-PEG hydrogels, 100 μL gels were formed containing the appropriate GelNB/PEG/LAP concentrations and either 50 nM Block-iT Oligonucleotide (Life Technologies) alone, or complexed with transfection agents. Gel mixtures were subsequently photo-polymerised and submerged in 100 μL PBS. Every 24 h, 5 μL PBS was removed and analysed using a Nanodrop (DeNovix DS-11 FX) to assess cumulative Block-iT oligonucleotide concentration in the supernatant.

### 2.7 Particle Size analysis

To determine the hydrodynamic size of DNA- (Block-iT fluorescent oligo) complexed transfection agents, each transfection agent was dispersed with targeted DNA at the respective concentration to achieve an N:P ratio of 5 in PBS, then subsequently incubated at RT °C for 30 min. Following which, the hydrodynamic size was assessed in triplicate using dynamic laser scattering (DSL) with a Malvern Zetasizer Nano ZS.

### 2.8 MSC Culture and Pre/Post Encapsulation miRNA Transfections

Human bone marrow derived MSCs (Lonza) were cultured in (DMEM containing D-glucose [1 g L^-1^] and sodium pyruvate [110 mg L^-1^] (Life Technologies), supplemented with penicillinstreptomycin [100 U mL^-1^] (Life Technologies) and 10 % (v/v) FBS. Cells were incubated at 37 °C with 5 % CO2 and passaged at 70 % confluency, reseeding was performed at a density of 2,500 cells cm^-2^. For our investigations, MSCs from 3 donors were used and all experiments were performed using cells at passage 6.

For *in situ* transfections, PEG, GelNB and LAP solutions were prepared as described previously. Cells were detached from culture dishes using TrypLE express (Life Technologies), resuspended into final hydrogel precursor solutions at a density of 2.5 million cell mL^-1^ and mixed thoroughly. Hydrogel-cell mixtures were pipetted into 30 μL drops on the bottom of 24-well culture dishes. Samples were then illuminated from above using visible light (400 – 500 nm, 10 mW cm^-2^) for 10 min, rinsed two times with 2 mL culture media, and maintained for the first 24 h. For basal culture conditions in which differentiation is not conducted, cells were maintained under culture conditions described above. For osteogenic induction, culture media was supplemented with 50 μM ascorbate-2-phosphate, 100 ng/mL dexamethasone and 10 mM -glycerophosphate 24 h after encapsulation. Media changes were performed every 4 days.

For miRNA transfections, human mercury LNA^*TM*^ miRNA mimics for miR-100-5p, miR-143-3p and mimic negative controls were used at concentrations of 10 nM, 25 nM and 25 nM respectively (**supplementary Table 2**). Transfections using polyethylenimine (PEI) (mw 4 kDa and 40 kDa (Sigma)) were performed using an N:P ratio of five. Lipofectamine RNA iMAX (Life Technologies) transfections were conducted as per the manufacturers protocol. For both *in situ* and pre-culture transfections, cells were serum-starved in 0.05 % FBS for 24 h prior to transfection. Pre-transfected cells were then incubated in transfection agents for a further 24 h prior to encapsulation. For *in situ* transfection conditions, all transfection reagents were combined with the GelNB/PEG/Cell suspension prior to photo-polymerisation.

### 2.9 MSC Viability Screening

MSC viability was assessed using a Live/Dead assay (Life Technologies) and CellTiter 96 proliferation assay kits (MTS) following the manufacturer’s instructions. Both assays were conducted at time points of 1, 3 and 7 days post-transfection. Live/Dead staining was imaged using a Nikon Eclipse TS2 inverted microscope (Nikon). CellTiter 96 proliferation assays were read using a Multiskan spectrum plate reader, using the SkanIT RE programme (version 2.4.2).

### 2.10 Flow cytometry Analysis

All flow cytometry analysis within this study was performed using a FACs Calibur flow cytometer (BD Biosciences). For 2D cultured experiments, cells were detached from the culture surface using TrypLE express (Life Technologies). For encapsulated scenarios, cell-laden hydrogels were incubated in Trypsin-EDTA (0.25 %) for 30 min at 37 °C until visible hydrogel degradation was observed, centrifuged at 1300 rpm for three min and resuspended in 1 mL [3 %] BSA in PBS (w/v) prior to analysis.

### 2.11 Quantitative real-time RT-PCR

To extract total RNA from encapsulated conditions, cell-laden hydrogels were incubated in Trypsin-EDTA (0.25 %) for 30 min at 37 °C until visible hydrogel degradation was observed and centrifuged at 1300 rpm for three min. Cells were subsequently washed three times in PBS, resuspended in 200 μL QIAzol Lysis Reagent. Total RNA was extracted using the Qiagen RNeasy mini kit with on-column DNase treatment (Cat #74104) according to the manufacturer’s instructions. Per condition 110 ng RNA was used for cDNA synthesis using Superscript VILO first strand synthesis master mix for qRT-PCR (Life Technologies) per manufacturer’s instructions in a total volume of 20 μL. Reverse transcription was performed in the T100 Thermal Cycler using the following cycling conditions: 10 min at 25 °C, 60 min at 42 °C and 5 min at 85 °C. Quantitative PCR reactions were set-up in a total volume of 10 μL with 1x ABI Fast SYBR Green Mastermix and 0.2 μM forward and reverse primers. The primer sequences are listed in **supplementary Table 1**. A CFX96 Real-Time System (Bio-Rad) was used to run the samples with fast cycling parameters of 20 s at 95 °C, 3 s at 95 °C and 30 s at 60 °C, which was repeated for 40 cycles and followed by a melt curve. Data was analysed by the 2^-ΔΔ^Ct method using RPS27a as a reference gene.

### 2.12 Staining and quantitation of mineral deposition

Osteogenic differentiation was quantified using xylenol orange (Sigma) and OsteoImage (Lonza) staining assays to detect mineral deposition. Quantitation was performed by normalising fluorescence intensity/unit area to cell number as determined by nuclear staining. To ensure systematic and non-biased analysis of the data sets, excitation wavelengths and powers were kept consistent between samples, with image acquisition conducted in identical regions.

For OsteoImage mineralization assays which detect hydroxyapatite deposition, samples were processed as per the manufacturer instructions. Briefly, cells were rinsed twice with PBS, fixed for 15 min in 4 % PFA diluted in PBS and stained with OsteoImage staining reagent [1:100] for 45 min. Cells were further washed thrice in PBS, permeabilized in PBS + 0.5 % Triton X-100 (Sigma) for 10 min, then incubated in blocking solution (PBS + 3 % BSA (Sigma)) containing Actin Readyprobe 555 (Life Technologies) for 30 min. Cells were subsequently washed in PBS five times, and imaged using a Nikon Eclipse Ts2 (Nikon) microscope.

For xylenol orange assays which detect calcium deposition, samples were washed with PBS and incubated in PBS containing Hoechst 33342 [1:1000 dilution] (Life Technologies) for 20 min. Samples were then rinsed thrice in PBS, followed by incubation in xylenol orange working solution (20 μM xylenol orange in dH_2_O) for 30 min. Samples were observed using a Nikon Eclipse Ts2 (Nikon) microscope. Mineral content stained with xylenol orange emits a red fluorescence which was then quantified using ImageJ to calculate corrected total cell fluorescence (CTCF) using the intensity density function. Background fluorescence was then subtracted from each data set, following which total fluorescence observed was divided by total cell number to provide a value representing mineral deposition per cell.

### 2.13 Statistical Analysis

Statistical analysis was conducted using GraphPad Prism 7 software, with at least three replicates across three independent MSC donors for each data point. Cell viability, gene expression and mineralisation measurements are displayed as the mean ± standard deviation, with one-way ANOVA and subsequent Tukey pairwise comparisons used to characterise statistical differences unless otherwise stated. Resulting p values of p ≤ 0.05 were deemed significant (*p ≤ 0.05, **p ≤ 0.005).

## 3 Results

### 3.1 Hydrogel characterisation and MSC encapsulation

A hydrogel system using thiol-ene photoclick reactions to couple norbornene-conjugated gelatin (GelNB) and PEG-dithiol (PEGdiSH) was selected as a base for our system. Previous data indicates high cell viability following encapsulation, together with the ability of these hydrogels to support MSC differentiation [18]. To further avoid the damaging effects of prolonged UV exposure, polymerisation was conducted outside of the UV range at 400-500 nm wavelengths, correlating to the tail end of the LAP activation spectrum. This optimised system has a high proportion of gelatin, which in turn facilitates cell-substrate interactions via integrin binding [19], thus making it highly suitable for our investigation.

By modulating GelNB concentrations, we were able to establish two hydrogels of distinct moduli, being either 4 % or 10 % (w/v) GelNB combined with 1 % (w/v) PEG-dithiol and 0.03 % (w/v) LAP (**Fig. 1a**), thus providing a versatile platform to assess roles of mechanical properties on miRNA delivery. Rheological characteristics of both compositions were assessed post-swelling to determine mechanical properties of the hydrogel present during prolonged incubation in cell culture medium. The 10 % GelNB compositions had G’ values of 680 ± 15 Pa, whereas 4 % GelNB hydrogels had a lower G’ value of 120 ± 20 Pa (**Fig. 1b**). Observations of swelling ratio revealed significant increases for 10 % GelNB hydrogels compared to those of 4 % counterparts, presenting values of 22.6 % ± 0.5 and 17.1 % ± 0.8 respectively (**Fig. 1c**). We also performed degradation studies to determine the suitability of the hydrogels to be used for the extended timeframe (weeks) required for MSC osteogenesis. The 4 % GelNB compositions began degrading after 15 days in culture, whereas the 10 % GelNB compositions were stable for up to 18 days before signs of degradation became evident (**Fig. 1c**).

**Figure 1:**
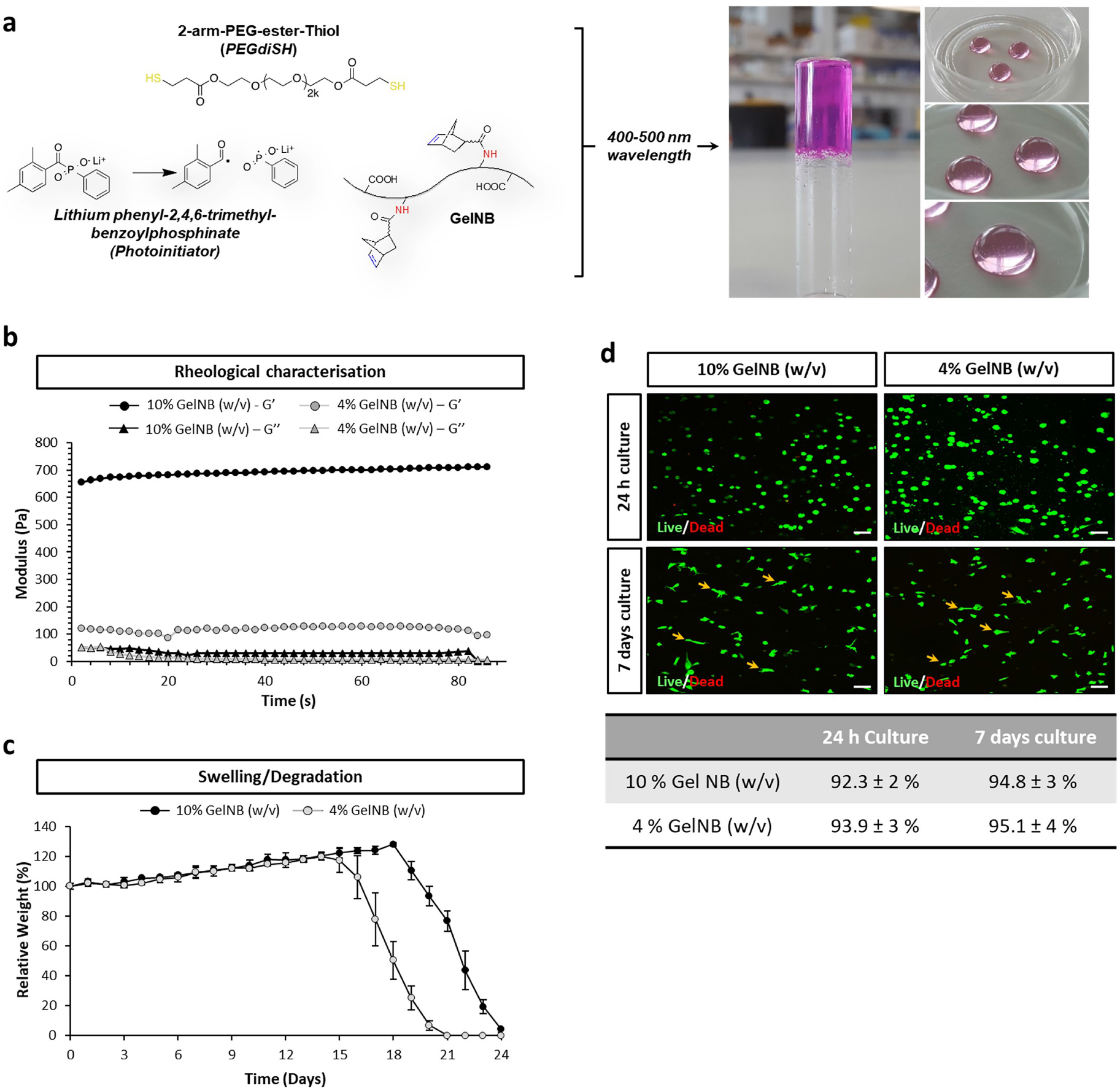
Characterisation of GelNB-PEG hydrogels. **a**. Structural composition of PEG-GelNB gels, and associated inversion test following photo-polymerisation under 400-500 nm wavelength light. **b**. Rheological characterisation of 10 % and 4 % (w/v) GelNB compositions post-swelling, showing storage (G’) and loss (G”) moduli. Data is presented as mean ± SD taken across three independent repeats, n = 12. **c**. Swelling and degradation characteristics of 10 % and 4 % (w/v) GelNB compositions incubated in cell culture media + FBS [10 %] at 37 °C. **d**. Representative Live/Dead (green/red) staining images of MSCs encapsulated in 10 % and 4 % (w/v) GelNB hydrogels under culture conditions. Arrows depict elongated cells. Tabulated values represent percentage of live cells from across three independent repeats (n ≥ 300 per condition). Scale bar, 50 μm.

To assess MSC viability, Live/Dead staining was performed at 24 h and 7 day timepoints. For both 4 % and 10 % GelNB compositions, high levels of cell viability were observed, with 92-94 % viability for 24 h cultures, and 94-95 % viability for 7 day cultures respectively (**Fig. 1d**). Good interaction of the cells with the hydrogel matrix are supported by observed changes in cell morphology, from spherical at 24 h to elongated phenotypes at 7 day. These findings not only suggest that a high proportion of the cells are viable both after initial encapsulation and extended culture, but that the cells can interact with the hydrogel matrix to adhere and spread.

### 3.2 Transfection of MSCs with miRNAs promotes osteogenesis in 2D

Having determined that the GelNB-PEG hydrogels could successfully support prolonged MSC culture, we next aimed to identify a suitable agent for miRNA transfection that had both high levels of efficiency and cell viability. The transfection agents tested were polyethylenimine (PEI) of two molecular weights (4 kDa and 40 kDa), previously demonstrated to facilitate cellular uptake of bound DNA through endocytosis as a result of charge interactions [20]. As a control, the commercially available Lipofectamine RNA iMAX was used. Although not clinically relevant, RNA iMAX has been extensively used as the gold-standard for miRNA and siRNA delivery and has previously shown to work well in MSCs [11].

Initial screening monitoring the cellular uptake of a fluorescently-tagged oligonucleotide sequence (Block-iT), revealed a high transfection efficiency for all agents, ranging from 97 % (RNA iMAX) to 77 % (4 kDa PEI) (**Fig. 2a**). Following successful transfection of MSC monolayer cultures, the transfection agents were screened for effects on both cellular viability and proliferation (**Fig. 2b** and **Fig. 2c**). Live/Dead staining showed no significant differences in MSC viability upon treatment with RNA iMAX or PEI at any timepoint. Furthermore, proliferation studies indicated no detrimental effects from any transfection agent, showing no significant difference in cell numbers across 24 h, 72 h or 7 days post-transfection.

**Figure 2:**
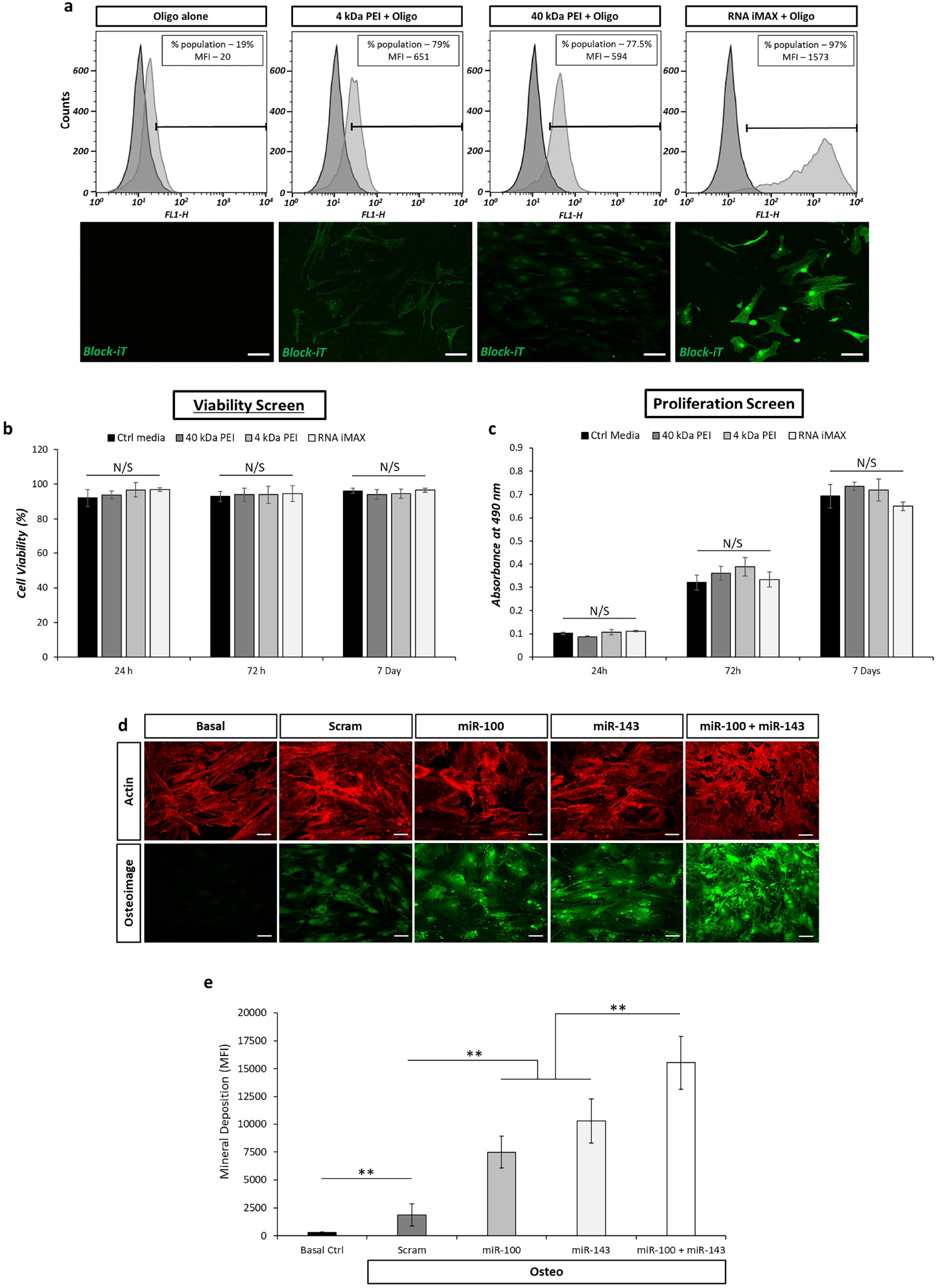
Transfection efficiency and miRNA effect on osteogenesis in 2D MSC cultures. **a**. Flow cytometry analysis and corresponding fluorescence staining demonstrate transfection agents ability to transport fluorescently tagged oligonucleotide sequences into MSCs following 24 h culture. Data presents percentage of positive cells within the total population together with observed mean fluorescence intensity (MFI). Fluorescence depicts the fluorescein tag from oligonucleotide sequences (green). Scale bar, 50 μm. **b**. and **c**. denote Live/Dead and MTS assay analysis of cellular viability and proliferation respectively across 1-, 3- and 7-day periods following 24 h agent incubation. All data is presented as mean ± SD for three independent MSC donors. Samples were analysed by one-way ANOVA with Tukey post hoc testing. **d**. Representative osteoImage fluorescence staining (green) counterstained with actin (red) depicting mineral deposition across miRNA supplemented MSCs cultured under osteogenic conditions. Scale bar, 50 μm. **e**. Quantification of mineral deposition per cell following either miR-100-5p, miR-143-3p or a combination of both miRNA mimics. Data is presented as mean ± SD taken across three independent repeats, n = 12. Samples were analysed by one-way ANOVA with Tukey post hoc testing. Statistically different samples are denoted by *p ≤ 0.05, **p ≤ 0.005

We next aimed to validate the pro-osteogenic effects of the miRNA mimics miR-100-5p and miR-143-3p, which we have previously demonstrated to enhance the osteogenic potential of MSCs in 3D cultures [11]. This time, 2D monolayer cultures were transfected using RNA iMAX, then cultured in osteogenic medium for 21 days, following which xylenol orange (for calcium) and OsteoImage (for hydroxyapatite) staining were used to determine the degree of mineralisation. As expected, MSCs cultured in basal medium did not form any mineral, whilst those cultured in the presence of osteogenic medium showed mineralisation to varying extents (**Fig. 2d**). MSCs transfected with miR-100-5p and miR-143-3p individually displayed increased mineral deposition compared to those transfected with a scrambled control miRNA. However, it was the samples transfected with both miR-100-5p and miR-143-3p that showed the highest coverage of mineralisation and most intense staining. Quantitative analysis (**Fig. 2e** and **Sup Fig. 1**) revealed these differences to be statistically significant, with a four-fold and three-fold increase in mineral deposition for individual miR-143 and miR-100 respectively. Mineralisation in samples treated with a combination of miR-100 and miR-143 was increased six-fold compared to scrambled controls. Together these data confirmed that this combination of miRNAs can be used to significantly enhance osteogenic differentiation of MSCs.

### 3.3 *In situ* transfection of MSCs encapsulated in gelatin-PEG hydrogels

We next incorporated miRNA:transfection agent complexes into the GelNB-PEG hydrogels and investigated both the release rate of these complexes and their ability to deliver miRNAs to encapsulated MSCs. Comparing both 10 % and 4 % GelNB compositions, transfection agents were incorporated directly into each hydrogel mixture prior to photo-polymerisation. The concentration of a fluorescently-tagged oligonucleotide (Block-iT) in the surrounding medium was measured over 7 days to evaluate release kinetics. For each gel composition, both Block-iT alone and transfection agent:Block-iT complexes were gradually released, rather than presenting a burst release response, with full release taking approximately 7 days (**Fig. 3a - 3d**). However, the release was consistently faster from the 4 % GelNB compositions compared to 10 % GelNB hydrogels, irrespective of the transfection agent. This is best exemplified under RNA iMAX-complexed conditions, in which half of the total oligonucleotide release from the 4 % GelNB hydrogels was observed at Day 4, whereas for the 10 % GelNB composition this value was not surpassed until Day 7.

**Figure 3:**
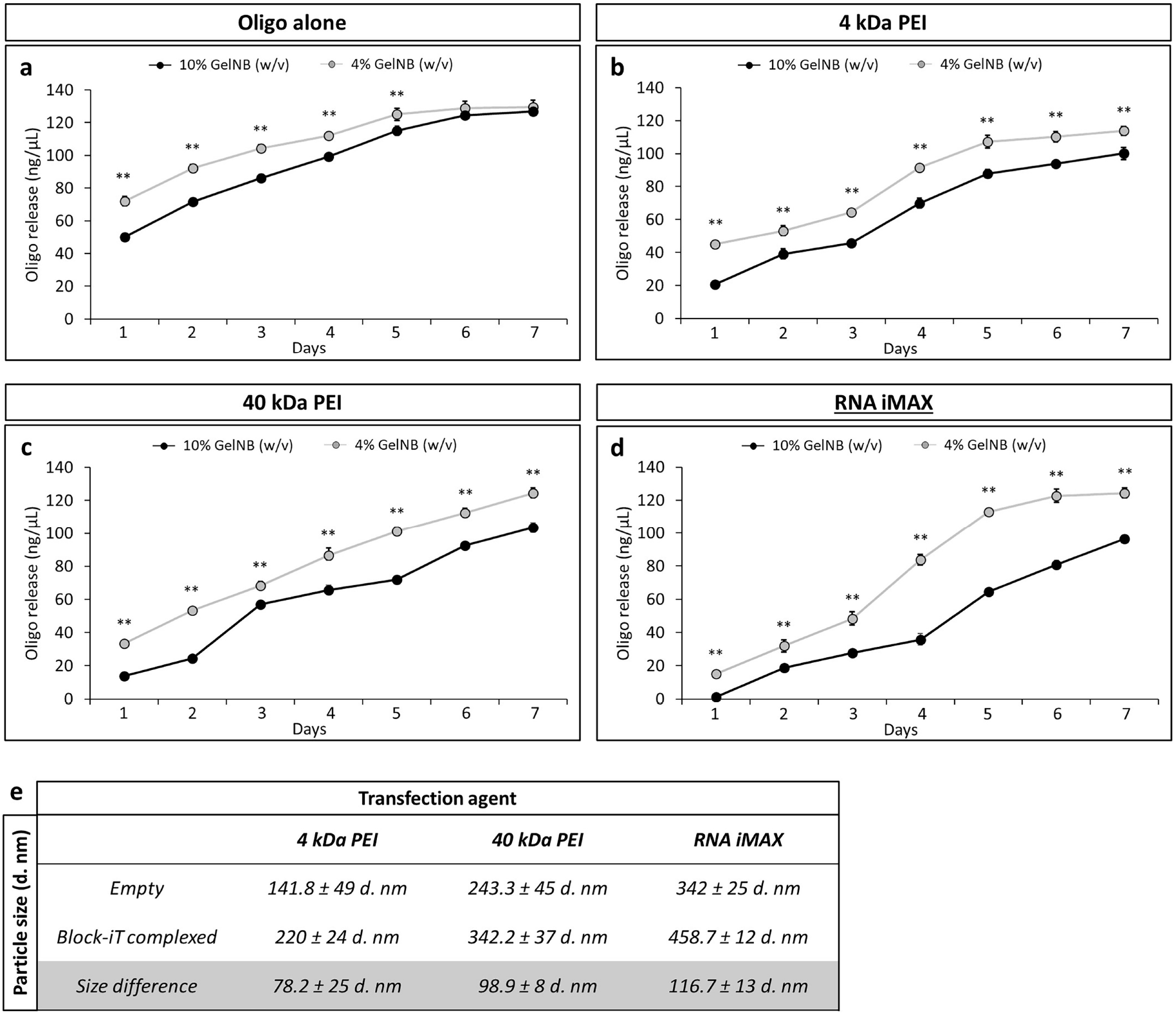
Kinetics of oligonucleotide and transfection agent complex release from 4 % and 10 % (w/v) GelNB hydrogels. Release kinetics of **a**. Uncomplexed oligonucleotide, **b**. 4 kDa PEI, **c**. 40 kDa PEI or **d**. RNA iMAX agents across 10 % or 4 % (w/v) GelNB hydrogels both individually and whilst bound to fluorescent oligonucleotides. Data is presented as mean ≤ SD for three independent repeats. Across each time point, comparisons between 10 % and 4 % GelNB compositions were analysed by t-test to discern statistical significance. **e**. Particle size analysis of chosen transfection agents either alone, or bound to fluorescent oligonucleotides. Tabulated data is presented as mean ± SD for three independent repeats, n = 24. Statistically different samples are denoted by *p ≤ 0.05, **p ≤ 0.005

To further probe these observations, hydrodynamic particle size analysis was performed on both empty and Block-iT-bound transfection agents (**Fig. 3e**). Interestingly, we observed consistent particle size increases for each transfection agent following Block-iT complexing (97 ± 19 d. nm), with the largest hydrodynamic particle size observed for complexed RNA iMAX (342 ± 25 d. nm), followed by 40 kDa PEI (243 ± 45 d. nm) and 4 kDa PEI (142 ± 49 d. nm) respectively. Comparing these data to the transfection agent release kinetics (**Fig. 3a - 3d** and **Sup fig. 2**), it is clear that the larger particles had slower release from their respective hydrogel compositions than smaller particles. For example, 40 kDa PEI and RNA iMAX complexed with Block-iT have the largest hydrodynamic radius and the least amount released from 10 % GelNB gels after 2 days (21 ng/ μL and 18 ng/μL respectively), whereas unbound oligonucleotide had the smallest hydrodynamic radius and a much greater amount released by day 2 (72 ng/μL). These findings indicate that transfection agent release is influenced by hydrogel composition and suggests that it may in future be possible to control miRNA delivery rate by modulating both hydrogel properties and transfection agent particle size.

We next examined whether it was possible to transfect encapsulated MSCs *in situ* from miRNA:transfection agent complexes incorporated within GelNB-PEG hydrogels. Transfection agent:Block-iT complexes and MSCs were incorporated into 10 % GelNB hydrogels and transfection efficiency determined after 72 h using immunofluorescence and flow cytometry to identify fluorescein-positive cells. The 10 % GelNB system was chosen due to its slower miRNA release compared to that of 4 % compositions, which we predicted would be more beneficial under extended culture conditions based on previous reports [21, 22]. MSCs were successfully transfected by all types of transfection agent. RNA iMAX presented the highest efficiency of 42 %, compared to 12 % and 18 % for the 4 kDa and 40 kDa PEIs respectively (**Fig. 4**). A comparison of these data shows that the transfection efficiencies were significantly lower than corresponding 2D transfections (**Fig. 2**), with each system displaying between 40 % - 60 % reduced overall efficiency. However, as both 2D and 3D experiments were conducted 72 h post encapsulation, where transfection complexes were incompletely released from the 3D scenario, we believe the decreased efficiency levels in 3D may partly be attributed to a lower amount of miRNA:transfection agent complexes being available to the cell at this point in time.

**Figure 4:**
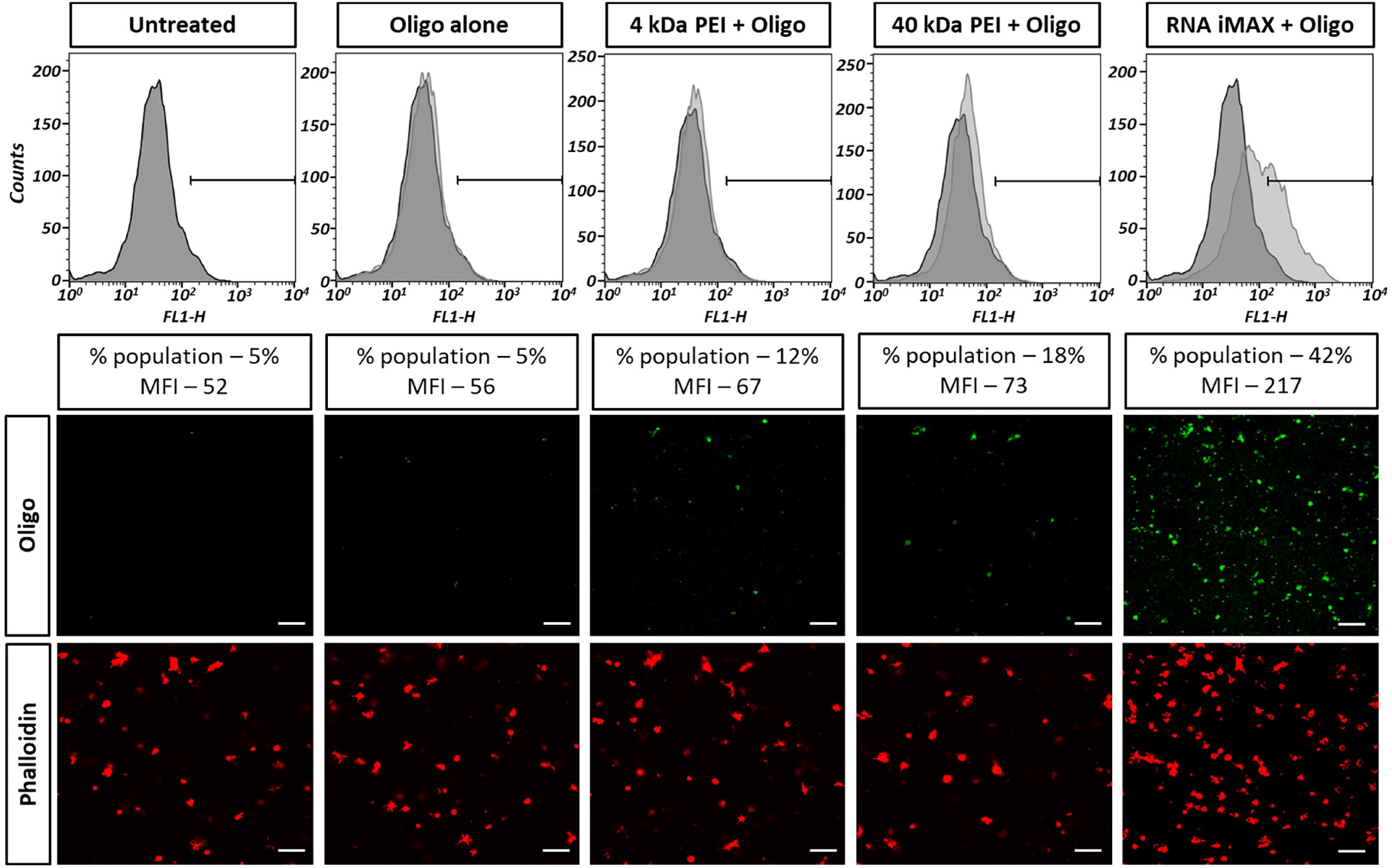
*In situ* transfection efficiency of MSCs encapsulated in GelNB-PEG hydrogels. Flow cytometry analysis and corresponding fluorescence staining observing chosen transfection agents ability to transport fluorescently tagged oligonucleotide sequences into MSCs *in situ* following 72 h culture. Data presents percentage of positive cells within the total population together with observed mean fluorescence intensity (MFI). Fluorescence depicts the fluorescein tag from the oligonucleotide sequence (green) and actin (red). Scale bar, 50 μm.

### 3.4 MSC encapsulation in miRNA loaded hydrogels increases osteogenic potential

To develop a simple one-pot, pro-osteogenic hydrogel system, we next tested the impact of delivering miR-100 and miR-143 directly to MSCs encapsulated in GelNB-PEG hydrogels. A comparison was made between *in situ* transfections using the different transfection agents, and also to MSCs that had been transfected 24 h prior to encapsulation (pre-transfected) using mineralisation as a marker of osteogenesis. For both pre-transfected and *in situ* transfected MSCs, an increase in mineral deposition was observed for all agents delivering miR-100 and miR-143, as compared to scrambled miRNA controls (**Fig.5**). Notably, for the *in situ* transfected samples, particularly intense mineral deposition was observed on the periphery of the hydrogel, irrespective of transfection agent used.

**Figure 5:**
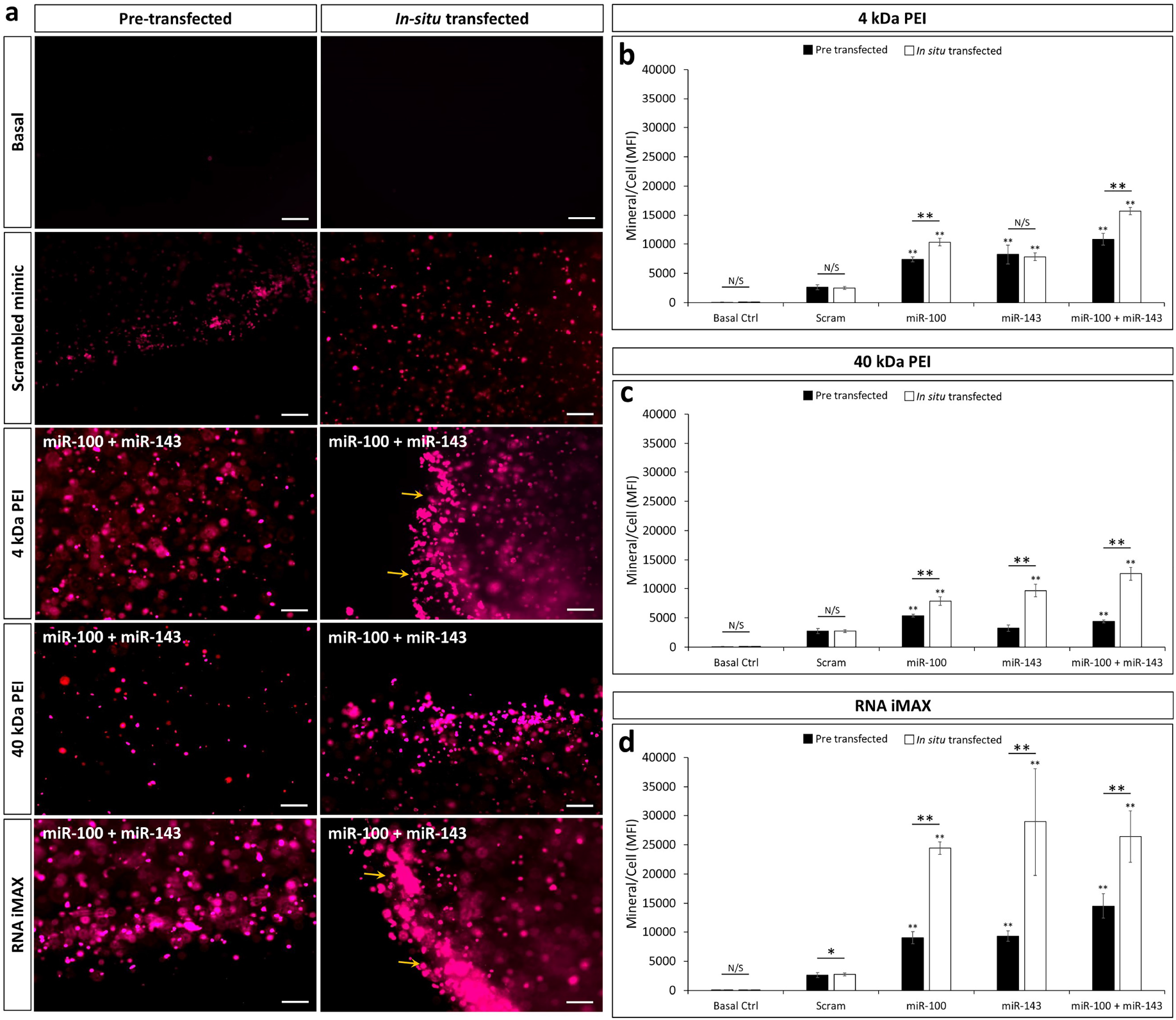
Comparison of mineralisation in pre-transfected vs. *in situ* transfected MSCs. **a**. Representative calcium staining using xylenol orange (red) across encapsulated MSCs either transfected prior to encapsulation or *in situ*. Staining depicts miR-100-5p and miR-143-3p combined transfection conditions using each chosen agent. Scale bar, 50 μm. **b. c**. and **d**. denote levels of calcium deposition per cell using 4 kDa PEI, 40 kDa PEI and RNA iMAX agents respectively following 21 d culture under osteogenic conditions. All data is presented as mean ± SD for three independent MSC donors, n = 60. Samples were analysed using unpaired t-tests and Welch’s corrections. Statistically different samples are denoted by *p ≤ 0.05, **p ≤ 0.005

Interestingly, further quantification revealed *in situ* transfections produced substantially more mineral than their respective pre-transfected condition in all but one combination (4 kDa PEI: miR-143-3p) (**Fig. 5b – 5d**). For 4 kDa PEI transfection agents, *in situ* transfections outperformed pre-transfected samples by 1.2-fold and 1.3-fold for miR-100 and miR-100/miR-143 respectively. For both 40 kDa PEI and RNA iMAX agents, the effect was even greater with mineralisation levels 1.5 – 2.2 times higher than their pre-transfected counterparts for all miRNAs tested.

To extend these findings, we analysed the expression of osteogenic marker genes under the different conditions (**Fig. 6**). Alkaline phosphatase expression was not significantly changed by pre-transfection of miR-100 or miR-143, but levels in the *in situ* transfected MSCs were significantly elevated, as compared both to MSCs transfected with a scrambled control miRNA and also to the equivalent pre-transfected cells (**Fig. 6a**). Similarly, whilst Col1a1 upregulation was only observed in RNA iMAX pre-transfections, all transfection agents caused upregulation of Col1a1 when delivered *in situ* (**Fig. 6b**). No significant differences were observed in the expression of osteopontin between transfection methods (**Fig. 6c**).

**Figure 6:**
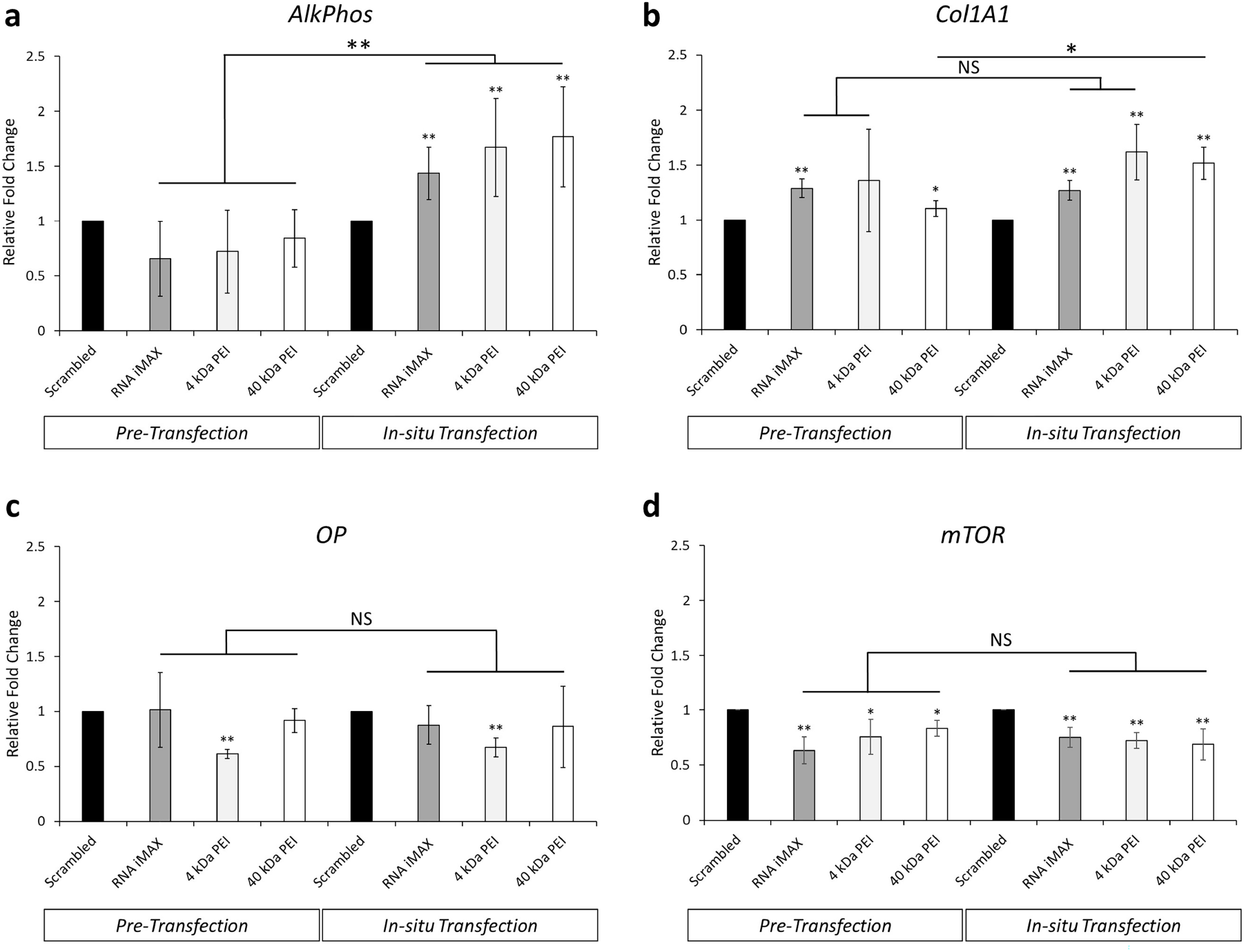
*in situ* transfection of encapsulated MSCs using combined miR-100-5p and miR-143-3p enhances osteogenic gene activity and decreases mTOR expression. **a. b. c**. and **d**. qPCR determination of the relative expression of osteogenic genes in MSCs treated with combined miR-100-5p and miR-143-3p mimics coupled to chosen transfection agents. All data is presented as mean ± SD for three independent MSC donors, n = 9. Samples were analysed by one-way ANOVA with Tukey post hoc testing. Statistically different samples are denoted by *p ≤ 0.05, **p ≤ 0.005

We have previously demonstrated that mTOR is a target of miR-100 and involved in the osteogenic effects of miR-100 via downregulation of mTOR [11]. Thus, whilst no significant differences were observed between the transfection agents or between pre- and *in situ*- transfected cells, levels of mTOR transcript were significantly downregulated in all samples transfected with miR-100 and miR-143, further confirming a role for mTOR in their regulation of osteogenesis (**Fig. 6d**).

## 4 Discussion

Our increasing understanding of mechanotransduction and its impact upon MSC fate provides enormous opportunity to more precisely guide MSC activities and differentiation. In particular, specific modulation of mechanotransductive signalling can be used to overcome limitations in the physical cues provided by biomaterials in tissue-engineering. One such example is the use of hydrogels for bone tissue-engineering. Hydrogels possess many properties that make them attractive for bone tissue engineering but are limited by the fact that their inherent softness does not provide the mechanical cues that promote osteogenesis. Here we have used mechanosensitive miRNAs known to enhance MSC osteogenesis to overcome this limitation of soft hydrogels and improve bone formation by MSCs in a GelNB-PEG hydrogel. miRNAs are attractive candidates for such an application due to their powerful regulatory effect on gene expression [16, 23]. We have previously demonstrated that miR-100-5p and miR-143-3p can promote MSC osteogenesis, but the future applicability of this approach was limited by the complexity of the system which required multiple stages of manipulation to transfect and then encapsulate the cells. In recent years, a handful of studies have probed the use of *in situ* hydrogel transfections for therapeutic application ranging from siRNA delivery *in vitro* to miRNA delivery *in vivo* [21, 24, 25, 26, 6, 27]. Interestingly however, these studies are limited to the use of either a single transfection agent or a limited range in mechanical properties for the base hydrogels. Here, we have developed a facile system that directly delivers miRNAs to encapsulated MSCs *in situ* using a range of transfection agents and shown that not only can this promote osteogenesis in a manner similar to pre-transfection, but that osteogenesis is actually enhanced.

As a base material for our system, we selected norbornene-conjugated gelatin (GelNB) and PEG-dithiol (PEGdiSH) to create our GelNB-PEG platform, which has previously been shown to support MSC viability, adhesion and differentiation [18, 28]. To further probe the effect of changing mechanical properties upon miRNA release and transfection rates, we modulated the weight percentage of GelNB within each hydrogel to create hydrogels of differing storage modulus. Interestingly, the values of storage modulus we obtained were lower than those previously reported [18] but we attribute this to the fact that our rheological measurements were obtained post-swelling in order to more accurately define the conditions presented to the encapsulated cells. This is in contrast to much of the field which conducts these measurements during the gelation process to monitor photo-polymerisation directly. Our degradation studies showed degradation within approximately 16 – 20 days for both 4 % and 10 % (w/v) GelNB compositions respectively. We attribute this relatively rapid degradation to hydrolysis of the ester groups within the thiolated PEG network [29, 30]. Nonetheless, the longevity of the hydrogels was sufficient for MSC differentiation and could therefore be used in our study. It is possible to slow the rate of degradation by removing the ester bond [28], however tissue-engineering aims to provide a scaffold to support new tissue formation with the goal of ultimately replacing the implant with endogenous tissue. For this reason, some degradation of the hydrogel is advantageous and could be modulated in the future by altering the ratio of ester and non-ester bonds to achieve a desired degradation rate.

We selected PEI as the transfection agent for our system due to both its ease of use and demonstrated ability to facilitate cellular uptake of bound DNA through endocytosis as a result of charge interactions [20]. We tested two distinct molecular weights (4 kDa and 40 kDa) due to the documented trade-off between transfection efficiency and agent toxicity, in which larger PEI polymers tend to display both the highest degree of efficiency and the highest degree of toxicity [31]. The commercially available, but not clinically relevant, RNA iMAX was used a control due to its well-documented efficacy and widespread use. Findings revealed no significant variation in transfection efficiency, metabolic activity or cell viability between each transfection agent in 2D transfections (**Fig. 2a, 2b and 2c**), suggesting that not only were the chosen agents able to deliver required oligonucleotide sequences, but were doing so with minimal impact to cellular function.

Our studies on the release rate of the transfection agents showed two clear trends. Firstly, release was more rapid from the 4 % GelNB compositions than the 10 % GelNB hydrogels. Secondly, release rate was correlated with the size of the complexes, as determined by particle size analysis. We attribute the faster release from the 4 % GelNB hydrogels to the lower cross-link density when compared to that of 10 % GelNB hydrogels. We hypothesise that the resulting decrease in mesh size would enable increased mobility of particles situated within the gels, allowing for faster diffusion out of the system. This premise can also be used to explain the differing release rate of particles of varying sizes as there was a general trend in that larger particles were released more slowly than smaller ones. These data suggest that by modulation of both transfection agent size and hydrogel composition, it may be possible to control transfection agent release speeds in the future, however further investigation is required to probe the potential influence of alternate factors such as charge-charge interactions between transfection complexes and the gelatin network.

Although consistent particle size increases were observed for 4 kDa PEI and 40 kDa PEI transfection agents following Block-iT coupling, the particle size increase for RNA iMAX agents were shown to exceed that of Block-iT addition alone. Similar observations have previously been reported by Ikonen *et al*, who suggested that as N/P ratios approach a zero zeta potential for transfection agent-DNA complexes (N/P 2.4 in this instance), polymer/DNA complexes begin to aggregate [32]. We believe that for our scenario, this exaggerated increase for RNA iMAX may represent the N/P ratios at which oligo/agent complexes migrate towards a neutral charge, and thus aggregate more readily with neighbouring particles.

Although it was possible to successfully transfect encapsulated MSCs directly from the hydrogel, a comparison of 2D and 3D transfections showed that rates were significantly reduced in the 3D system (**Fig. 4**). One reason for the lower efficiency may be that the *in situ* transfection tests were conducted after 72 h culture which, when compared to the timescale of transfection agent release (**Fig. 3**) is at the point when we observe approximately 20 %, 40 % and 50 % oligonucleotide release for RNA iMAX, 4 kDa PEI and 40 kDa PEI agents respectively. However, the final impact upon osteogenesis suggests that the influence on MSC fate is significant and actually enhanced for *in situ* transfection. This confirms successful delivery of the miRNAs and may indicate that sustained contact with transfection agents over a prolonged period of time is better than a short, high-concentration treatment, as in the 2D methodology. However, further studies would be required to fully determine the impact of transfection agent contact on transfection rates and subsequent changes in target gene expression.

miR-100-5p and miR143-3p have been shown to increase mineralisation in 3D (via pre-transfected cells), but this phenomenon has never been tested in 2D or *in situ* transfections. Our results validate this combination of miRNAs, in particular confirming dual transfection of both candidates as a particularly powerful way to increase osteogenesis. Although both miR-100 and miR-143 converge upon the mTOR signalling network, the boost observed when combining the miRNAs may be because these have distinct mRNA targets, with miR-100 targeting mTOR itself and miR-143 targeting RICTOR and LARP [11, 33, 34, 35]. Regulation of osteogenesis via the mTOR pathway in this study was further validated by the reduced mTOR transcript level in all of the transfected MSC populations (**Fig. 6d**). When coupled with evidence suggesting direct modulation of mTOR is able to regulate alkaline phosphatase activity after only 24 h induction [36], we suggest that as our select miRNAs target mTOR signalling, our findings may be a result of alternate alkaline phosphatase modulation than that of osteogenic differentiation alone, however further investigation will be needed to validate this.

Following the optimisation of each individual component, we proceeded to validate the ability of our combined *in situ* transfection platform to promote osteogenic potential of MSCs. The utility of miR-100 and miR-143 was again validated by their ability to significantly enhance mineralisation under all conditions tested, with the exception of 4 kDa PEI:miR-143-3p (**Fig 5**). This observation may be attributed to 4 kDa PEI transfections having the lowest efficiency and therefore sub-optimal effects on osteogenesis (**Fig. 4**).

Unexpectedly, the *in situ* transfections consistently outperformed the pre-transfected samples, both in terms of osteogenic gene expression and mineralisation. This suggests that the *in situ* transfection of miRNAs may not only be a simpler system to use but also a more effective one. The mechanism behind this is unclear, but one possible explanation is that increasing the duration of exposure of MSCs to the transfection agents may lead to improved outcomes. Support for this theory comes from data presented by Zhang *et al*, in which sustained release of miR-21 over a 2 week period enhanced target gene modulation to that of bulk released scenarios [37]. Assessment of gene regulation further supported that through *in situ* transfection, successful modulation of pro-osteogenic genes was observed (**Fig 6**). Although comparable increases in Col1A1 expression between pre- and *in situ* transfection conditions are present (**Fig. 6b**), we did observe variation for 40 kDa PEI agents, revealing a significant increase in gene activity for *in situ* transfection, which we attribute to the decreased diffusion rates observed for *in situ* release. We suggest that these decreased diffusion rates may increase contact time between miRNA loaded agents and MSCs, resulting in increased activation of Col1A1 to that of pre-transfected conditions which were subject to only a 24 h contact time.

## 5 Conclusion

We have developed and tested a facile system for *in situ* miRNA transfection and demonstrated its utility to promote osteogenesis of MSCs in a soft hydrogel. The addition of miR-100-5p and miR-143-3p, when complexed with PEI for transfection, provides a simple, yet effective platform to maintain a gradual release of target miRNAs to encapsulated cells. *In situ* transfection was more effective than encapsulation of pre-transfected MSCs, despite a lower apparent transfection efficiency at 72 h. Our data also suggest that there is scope to further tailor the release rate and consequently the timing or efficiency of *in situ* transfections by modifying base hydrogel formulation and/or transfection agent. We believe that our strategy, which combines modulators of mechanotransduction with a simple *in situ* transfection system, provides a promising means to improve bone formation by MSCs in hydrogels, but may also hold future utility in tissue-engineering across a much broader range of cell and tissue types.

## 6 Supplementary Information

**Supplementary Figure 1:**
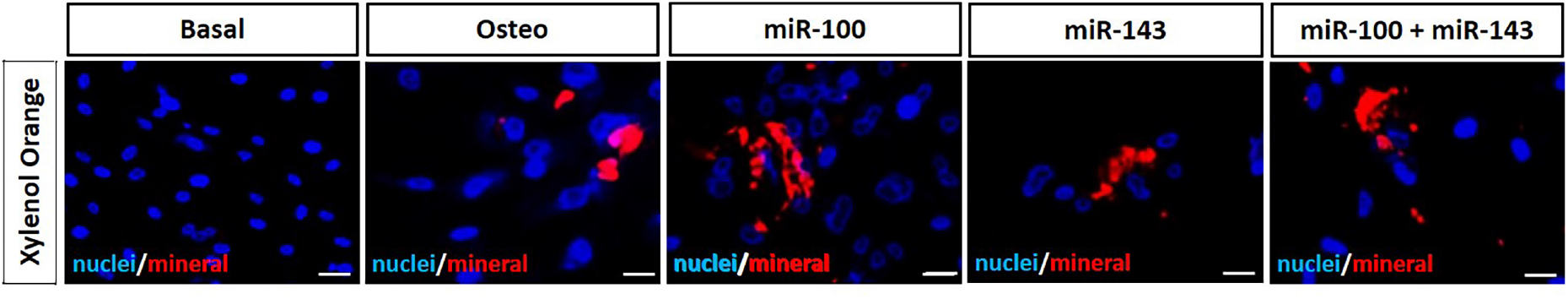
microRNA addition stimulates increased calcium deposition per cell. Comparative calcium deposition staining of 2D cultured MSCs transfected with screened microRNAs using RNA iMAX. Images depict nuclei (blue), and calcium (red), enabling quantification of mineral/cell to be calculated. Scale bar, 10 μm.

**Supplementary Figure 2:**
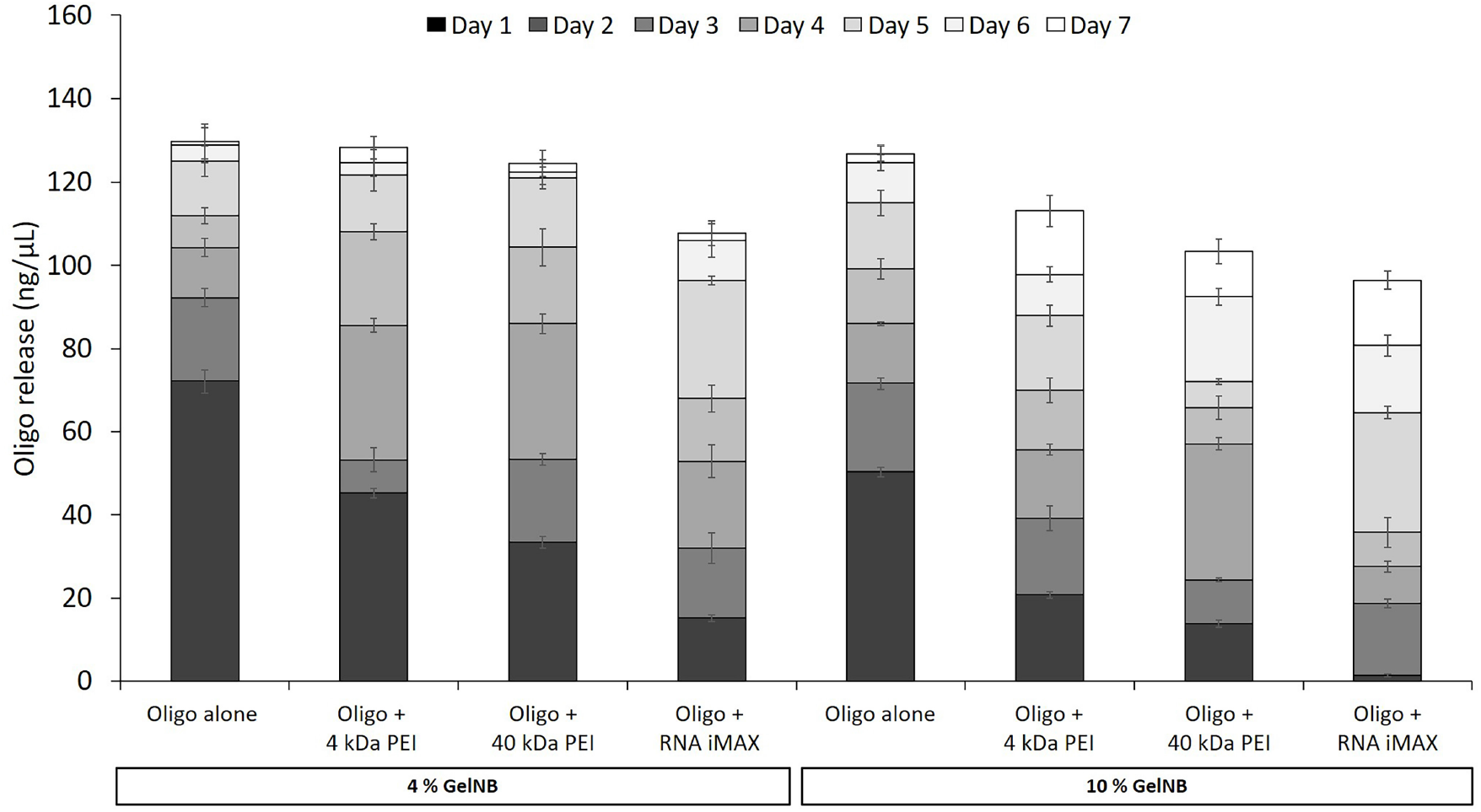
Release kinetics of oligo:transfection agent complexes from 4 % and 10 % GelNB hydrogels. Daily observation of oligonucleotide release from 10 % and 4 % (w/v) GelNB hydrogel compositions between chosen transfection agents. Daily data is presented as mean ± SD from three independent repeats, n = 9.

**Supplementary Figure 3:**
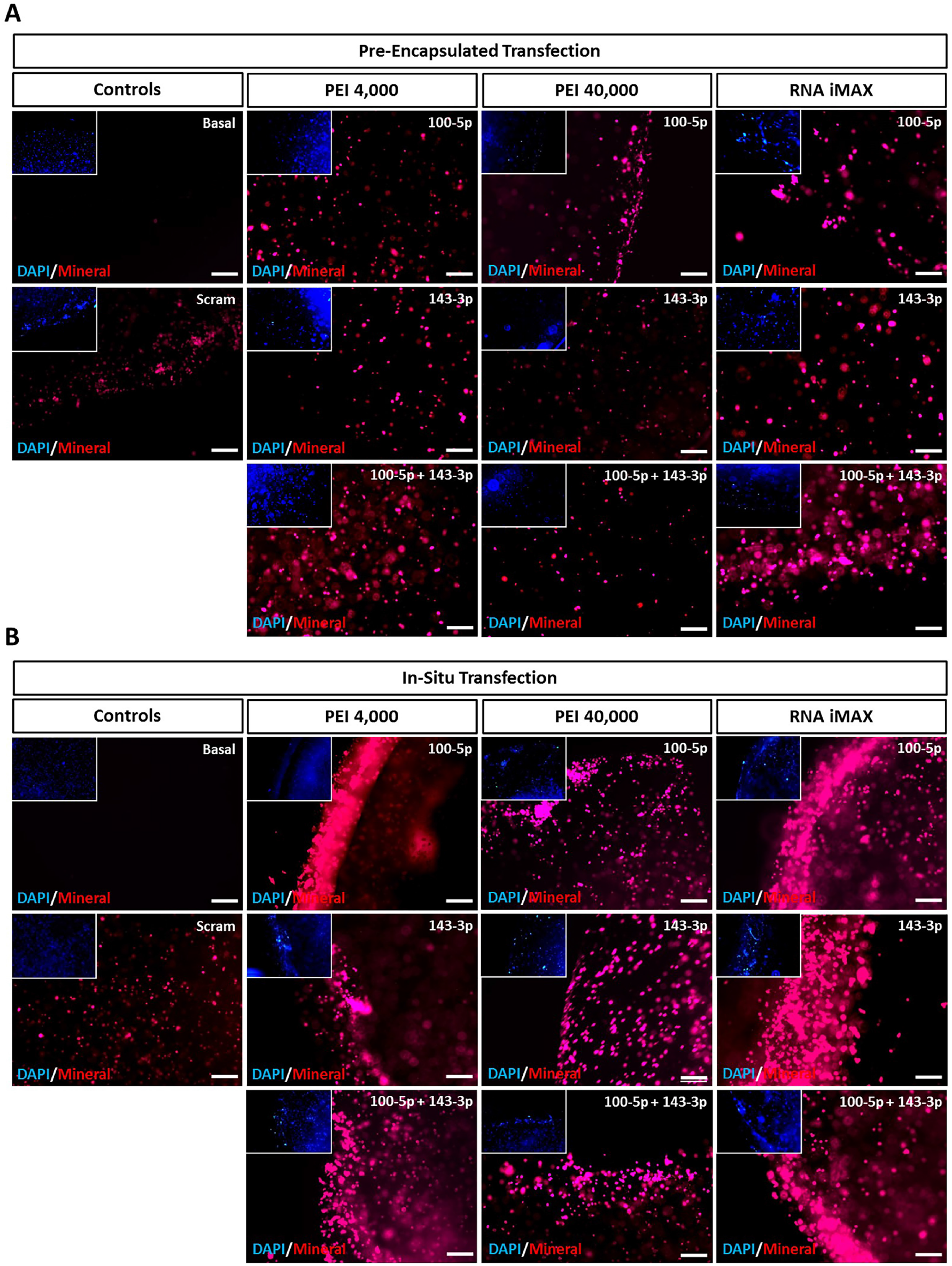
Mineral deposition by *in situ* transfection of MSCs encapsulated in 10 % (w/v) GelNB hydrogels together with a miR-100-5p, miR-143-3p. **a**. and **b**. Depict calcium deposition (red) and nuclear staining (blue) for MSCs either pre-transfected prior to encapsulation or transfected *in situ* following 21 days culture under osteogenic conditions. Representative images for each chosen transfection agent/miRNA condition are displayed. Scale bar, 50 μm.

**Supplementary Figure 4:**
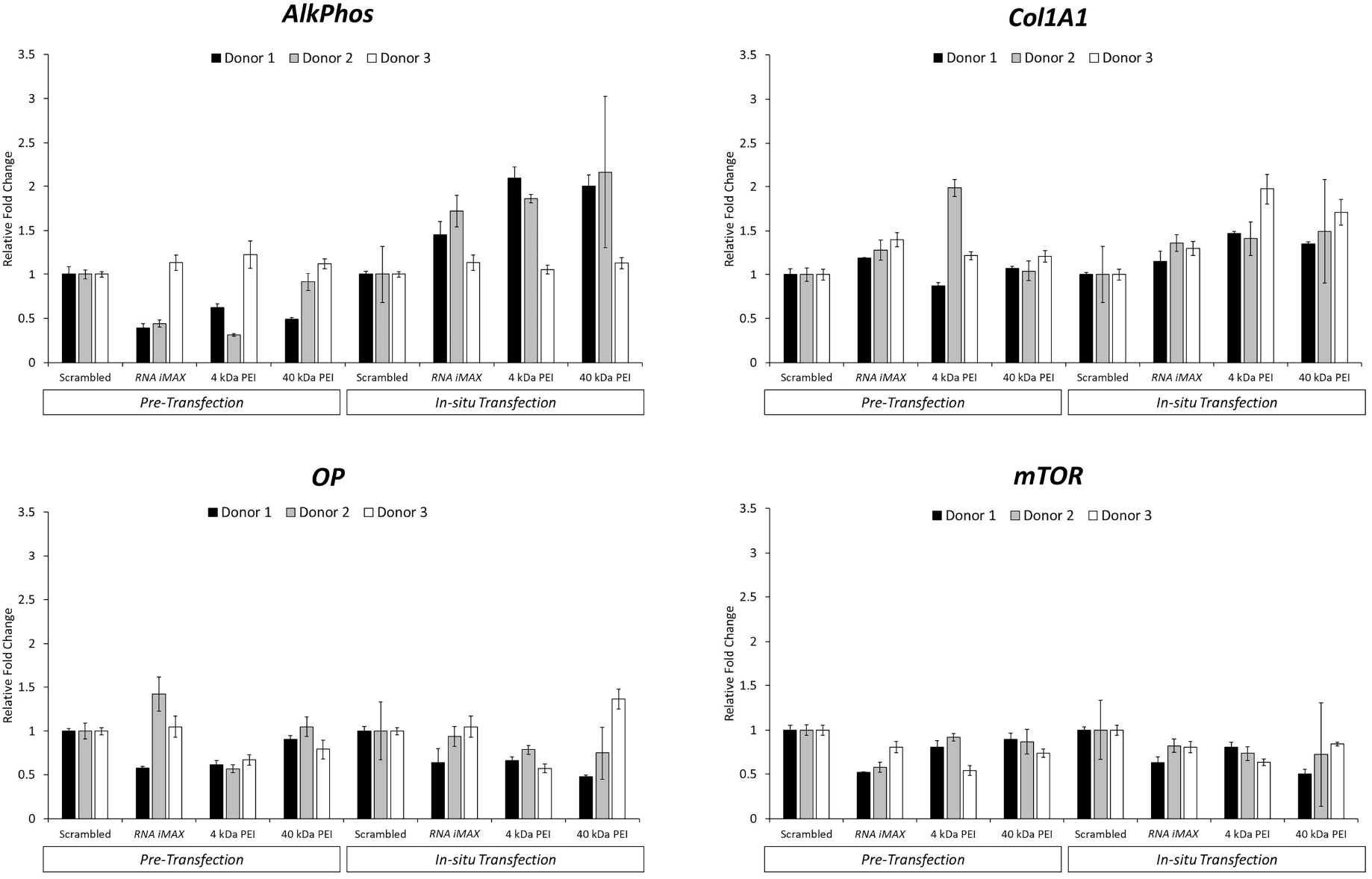
*in situ* transfection of encapsulated MSCs using combined miR-100-5p and miR-143-3p enhances osteogenic gene activity and targets mTOR expression. Individual donor qPCR data depicting relative expression of osteogenic genes in MSCs treated with combined miR-100-5p and miR-143-3p mimics coupled to chosen transfection agents. All data is presented as mean ± SD, n = 3.

**Supplementary Table 1:**
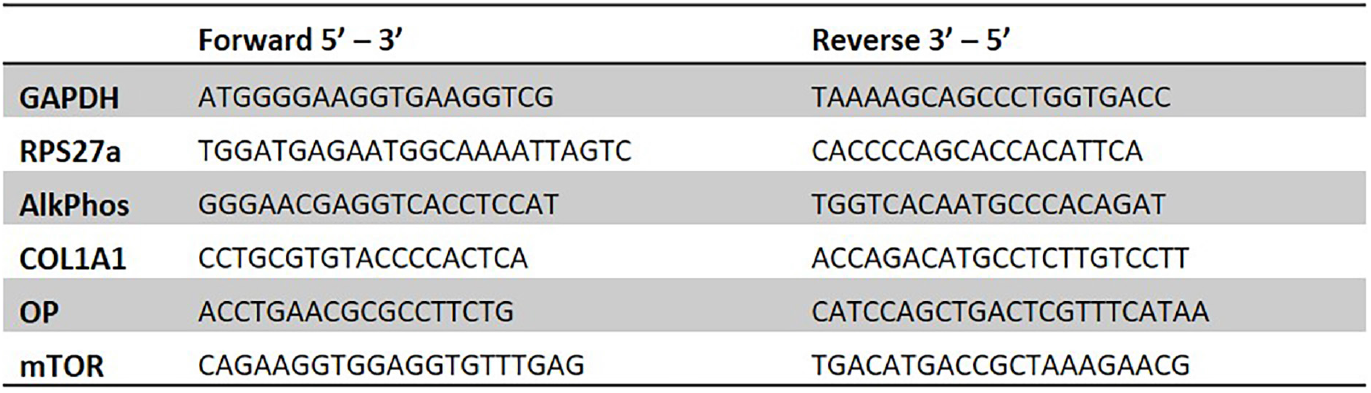
qPCR forward and reverse primer sequences used in this study. GAPDH: glyceraldehyde-3-phosphate dehydrogenase. RPS27a: ribosomal protein S27a. AlkPhos: alkaline phosphatase. COL1A1: collagen type I alpha 1 chain. OP: osteopontin. mTOR: mammalian target of rapamycin.

**Supplementary Table 2:** Exiqon miRNA mimics used in this study.

|  | miRNA mimic |
| --- | --- |
| **Mimic negative control, FAM labelled** | 479995-011 |
| **hsa-miR-100-5p** | 472477-001 |
| **has-miR-143-3p** | 470035-001 |

